# Serum hormone levels in infertile women between conceived with and without hormone replacement therapy: Safety of estradiol and progesterone replacement therapy in frozen-thawed embryo transfer cycles for breast cancer survivors

**DOI:** 10.1101/2021.01.29.428765

**Authors:** Ayumu Ito, Yukiko Katagiri, Yusuke Fukuda, Mineto Morita

**Affiliations:** Department of Obstetrics and Gynecology, Toho University Graduate School of Medicine, Ota-ku, Tokyo, Japan; Department of Obstetrics and Gynecology, Faculty of Medicine, Toho University, Ota-ku, Tokyo, Japan; Reproduction Center, Toho University Omori Medical Center, Ota-ku, Tokyo, Japan

## Abstract

**Objective:** Recent advances in cancer treatment and reproductive medicine have made the post-treatment quality of life an important concern for cancer survivors. We aimed to evaluate the safety of sex hormone (estradiol and progesterone) replacement therapy (HRT) in women who conceived by assisted reproductive technology (ART) with hormone receptor-positive breast cancer.

**Methods:** We measured serum E2 and P4 levels at 4–10 weeks of gestation in women who conceived naturally or after timed intercourse or intrauterine insemination for infertility without HRT for luteal support (non-HR group; n=135). We conducted a retrospective comparison of the values from the non-HR group with those of women who conceived by ART with HRT for infertility (HR group; n=75).

**Results:** Serum E2 levels were significantly higher in the non-HR group than in the HR group at 5, 6, and 8 weeks of gestation. Similarly, serum P4 levels were significantly higher in the non-HR group than in the HR group at 4, 5, and 6 weeks of gestation.

**Conclusions:** This study suggests that in cancer reproductive medicine for hormone-dependent breast cancer survivors, HRT administered during the first trimester of a pregnancy after primary disease treatment may not increase the sex hormone levels to levels above those seen in spontaneous pregnancy.

## Introduction

Recent advancement in cancer treatment and reproductive medicine has increased the importance of post-treatment quality of life among childhood, adolescent, and young adulthood cancer survivors. Fertility preservation is of major concern, and the use of assisted reproductive technology (ART), such as cryopreservation of embryos, oocytes, or ovarian tissue, is important for conserving fertility. For women with a history of breast cancer and other hormone-sensitive malignancies, hormone replacement therapy (HRT), which is important for continued pregnancy using ART, risks the exacerbation or recurrence of the primary disease. Hormonal exposure owing to pregnancy could be also a risk factor; however, some studies have reported that the prognosis of breast cancer survivors who underwent appropriate neoadjuvant therapy is not necessarily worsened by spontaneous pregnancy [1] [2] [3] [4] [5]. Although the rates of recurrence and mortality are lower for patients who receive multidrug chemotherapy following breast cancer surgery than for those treated by surgery alone [6], patients who receive multidrug chemotherapy show decreased fertility owing to chemotherapy-induced ovarian failure and age-associated ovarian dysfunction resulting from prolonged administration of hormone therapy. Furthermore, many patients encounter difficulty in conceiving naturally. The levels of anti-Mullerian hormone (AMH), a parameter of ovarian reserve, decrease to undetectable levels during chemotherapy and remain low even after completing of chemotherapy [7]. Studies have reported low rates of pregnancy in breast cancer survivors [8] [9] [10]; hence, before initiating treatment, patients must be provided with adequate information, informed consent should be obtained, and patients should be offered consulting with a doctor specializing in reproductive medicine. Under these circumstances, the standard practice is to offer various options for preserving fertility such as cryopreservation of embryo, oocytes, or ovarian tissue to women with cancer in the limited duration between their diagnosis and treatment. However, the benefits and drawbacks of ovarian stimulation, embryo transfer, and HRT as part of ART are unclear with respect to the effect on the primary disease and are currently a subject of debate. In particular, if ovarian function decreased and ovulation is impaired after cancer treatment, HRT is essential for embryo transfer using frozen embryos or oocytes once the patient is allowed to conceive. Even after pregnancy is established, this support must be continued until the main site of hormone production switches from the corpus luteum to the placenta (the luteo-placental shift).

In this study, we compared the hormone levels during the first trimester of pregnancy between women who conceived naturally or after timed intercourse (TI) or intrauterine insemination (IUI), which does not require HRT for infertility, and those who conceived with ART. We aimed to evaluate the safety of estrogen and progesterone replacement therapy for frozen-thawed embryo transfer in estrogen and progestogen replacement therapy for frozen-thawed embryo transfer in women with hormone receptor-positive breast cancer who conceived using ART to compare with those who conceived naturally or after TI or IUI.

## Materials and Methods

We measured the serum E2 and P4 levels at 4–10 weeks of gestation in non-HR group participants. The study subjects were women treated in the Department of Obstetrics and Gynecology or Reproduction Center, Toho University Omori Medical Center, between November 2018 and April 2019, who conceived naturally or after TI or IUI for infertility. The non-HR group participants did not undergo HRT for luteal support (Fig 1). TI and IUI cycles were performed naturally or using medication for ovulation induction with follicle growth monitoring by ultrasonography (Fig 2).

**Fig 1.**
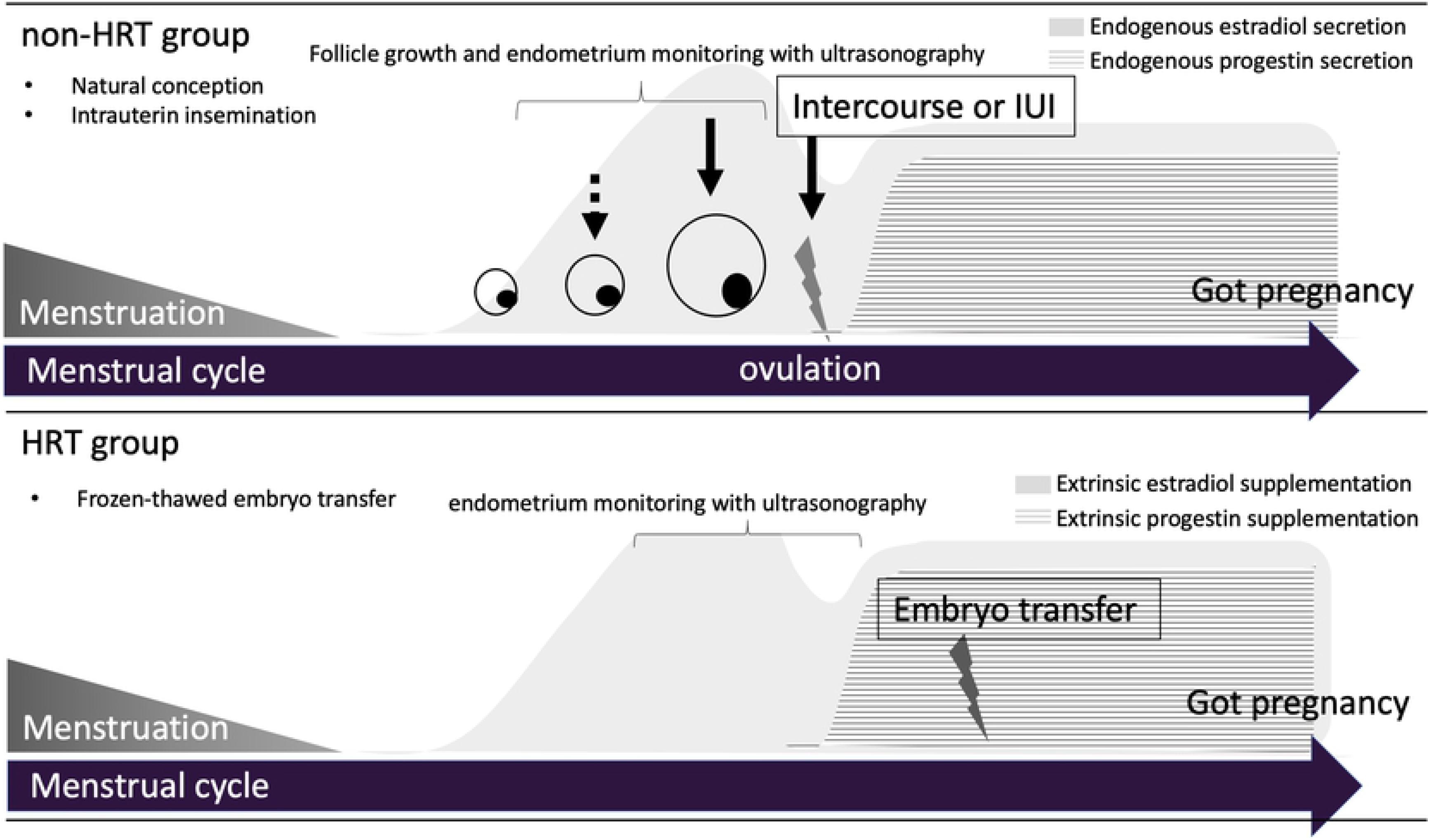
Medical treatment schedules of non-HR group and HR-group

**Fig 2.**
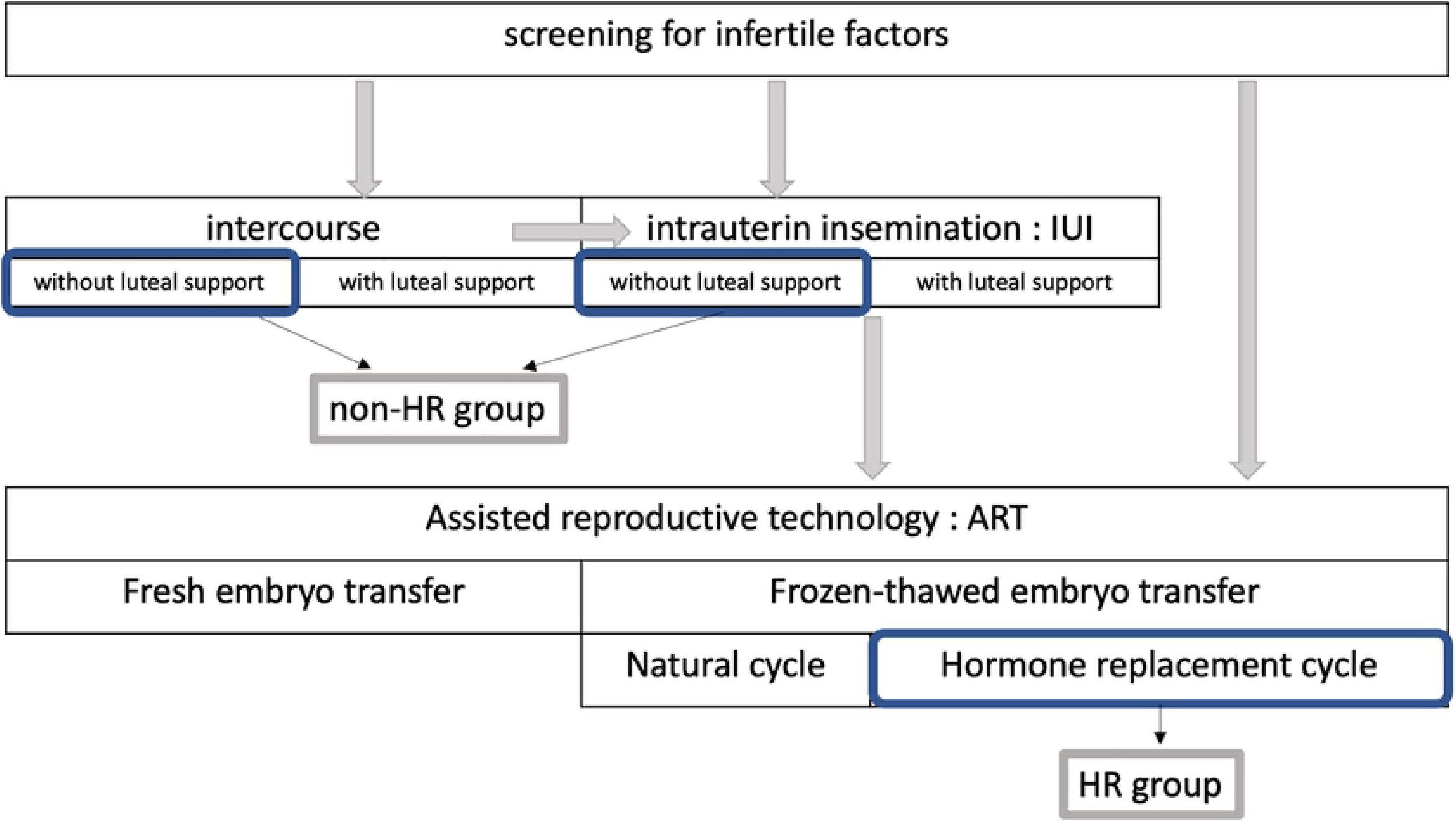
Flowchart of fertility treatment

We retrospectively compared the serum E2 and P4 levels at 4–10 weeks of gestation between the non-HR and HR groups. The HR group included women who conceived after frozen-thawed embryo transfer and HRT with estrogen and progesterone in our reproduction center between January and December 2018 (Fig 1). The members of both groups continued their pregnancies until at least 12 weeks of gestation.

Estrogen replacement therapy was administered in the form of transdermal estrogen patch applied every alternate day from day 3 of menstruation. The initial dose was 2.16 mg, and this was increased after a few days to 2.88 mg and then to 3.60 mg. Transvaginal natural progesterone at a dose of 90–800 mg/day was administered when the thickness of the endometrium was ≥8 mm; depending on the stage of the frozen embryo to be transferred, the embryo transfer was performed on day 2 (P+2), day 3 (P+3), or day 5 (P+5) after the first day of administration of transvaginal natural progesterone use (P0). The serum hCG level was measured at 4w0d of gestation counted from the date of transfer. If the result was positive, HR was continued, and pregnancy was confirmed when the gestational sac was visible. The serum E2 and P4 levels were subsequently measured, and the administration of transdermal estrogen and transvaginal natural progesterone was continued until adequate endogenous hormone secretion was confirmed (Fig 3).

**Fig 3.**
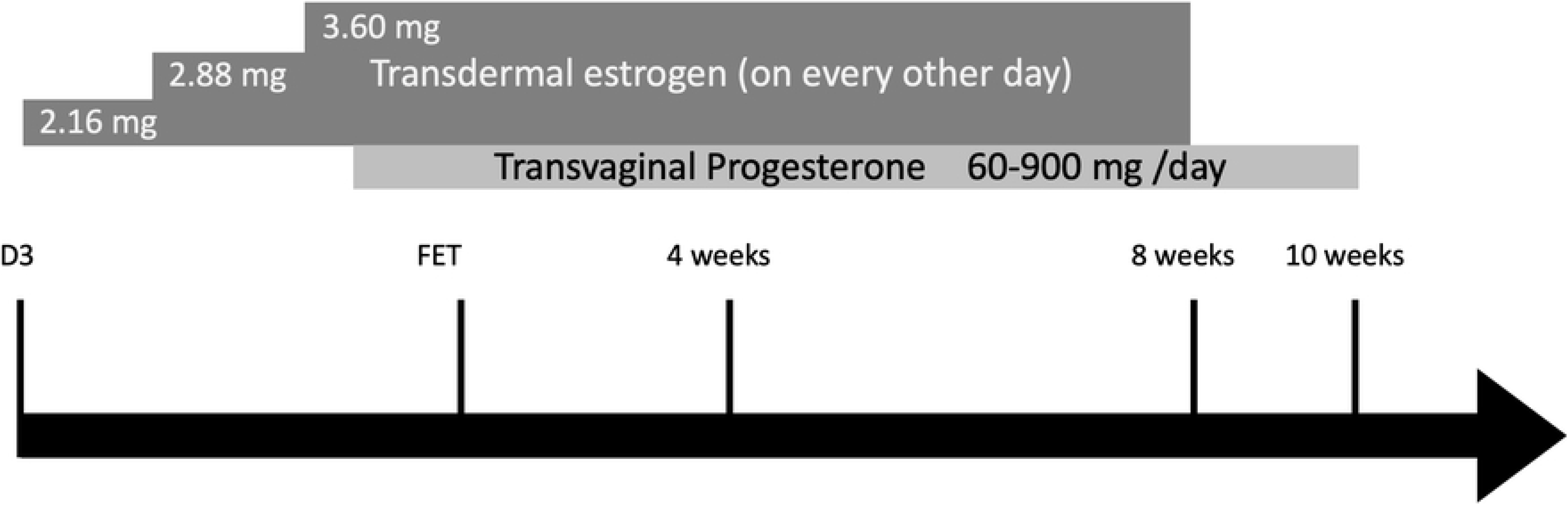
Basic protocol for frozen-thawed embryo transfer in a hormone-induced ovulatory cycle and luteal support. Estrogen replacement therapy was administered in the form of transdermal estrogen patch applied every alternate day from Day 3 of menstruation. Transvaginal natural progesterone was administered when the thickness of the endometrium was ≥8 mm; depending on the stage of the frozen embryo to be transferred, the embryo transfer was performed.

SPSS Statistics ver. 25 (SPSS Inc., Chicago, IL, USA) was used for statistical analysis.

### Ethics declarations

The protocol of the study was approved by the Ethics Committee of Toho University Omori Medical Center (Approval No. M1704717209 and M18239) and conformed with the 1964 guidelines of the Helsinki Declaration and its later amendments. Informed consent was obtained from all patients. We obtained written informed consent was obtained from the patients of non-HR group and informed consent in the form of opt-out on the website from the patients of HR group. Information on the research was made public on the website of each institution, and the opportunity for the research subjects to refuse participation was guaranteed.

## Results

The study population included 135 women who conceived naturally without the use of HRT (non-HR group; of these, 100 had conceived naturally and 35 had conceived after IUI) and 75 who conceived after frozen-thawed embryo transfer and HRT with estrogen and progesterone in our Reproduction Center between January and December 2018 (HR group). There was no significant difference in the patient age between the two groups (33.4±5.0 years vs. 35.3±4.2 years, non-HR group vs. HR group).

The serum E2 and P4 levels in the non-HR and HR groups increased over time during the first trimester of pregnancy (Figs 4a, 4b, 5a, and 5b). There was no significant difference in the pregnancy continuation rate between the two groups. The formulas for the approximate curves of the serum E2 and P4 levels for the non-HR group are y=234 x+ 500 (R^2^=0.17) and y=0.8x + 23.3 (R^2^=0.027) and those for the HR group are y=258x−112 (R^2^=0.63) and y=2.5x + 9.3 (R^2^=0.27).

**Fig 4.**
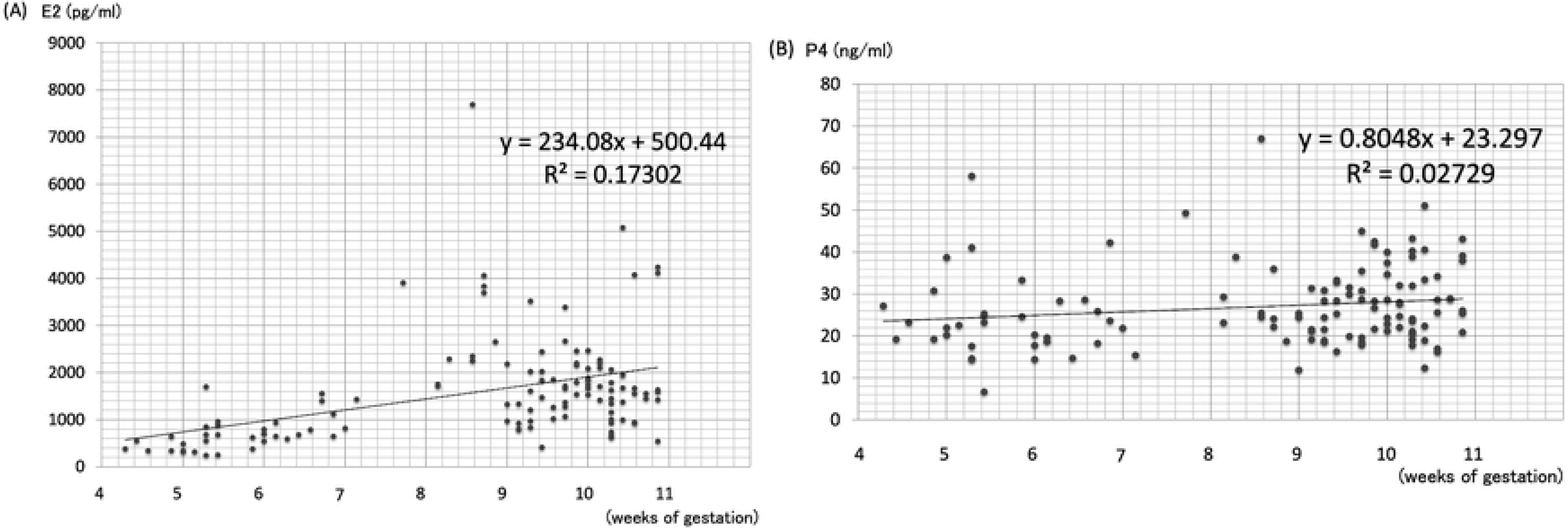
Serum hormone levels in the non-HR group. (a) Serum E2 levels during the first trimester in the non-HR group. (b) Serum P4 levels during the first trimester in the non-HR group.

**Fig 5.**
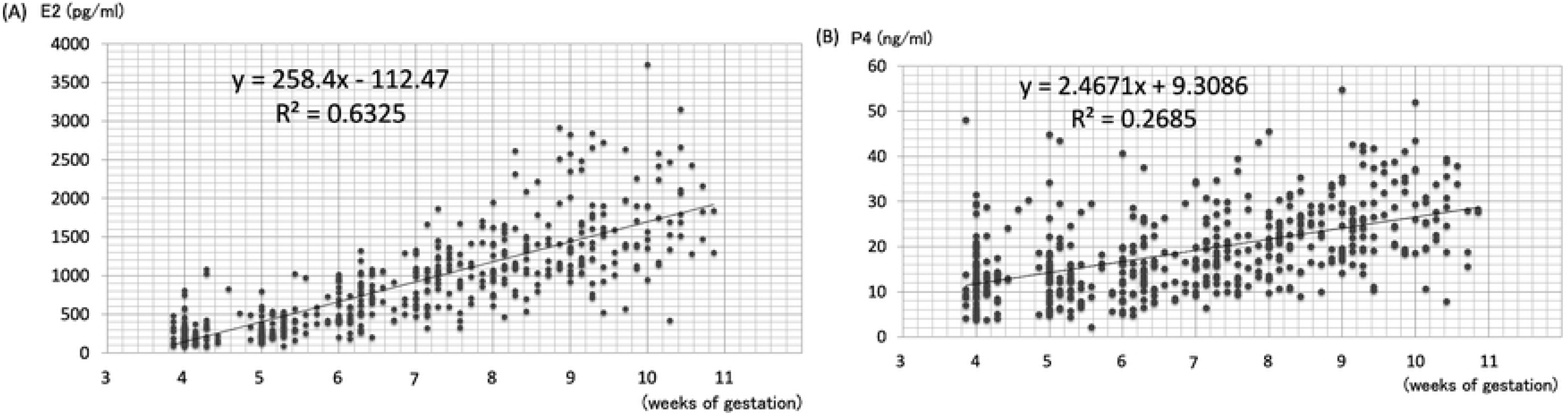
Serum hormone levels in the HR group. (a) Serum E2 levels during the first trimester in the HR group. (b) Serum P4 levels during the first trimester in the HR group.

The serum E2 values in the non-HR and HR groups were 453.8±137.7 and 291.9±200.7 pg/mL at 4 weeks, 622.3±379.7 and 351.3±173.0 pg/mL at 5 weeks, 870.1±326.7 and 644.1±262.3 pg/mL at 6 weeks, 2056.3±1635.0 and 976.6±342.2 pg/mL at 7 weeks, 3232.0±1781.3 and 1285.7±453.4 pg/mL at 8 weeks, 1589.3±660.9 and 1556.3±587.9 pg/mL at 9 weeks, and 81790.9±974.6 and 1781±649.2 pg/mL at 10 weeks of gestation, respectively. The serum E2 level was significantly higher in the non-HR group than in the HR group at 5, 6, and 8 weeks of gestation (5 weeks, p <0.05; 6 weeks, p <0.01; 8 weeks, p <0.01). There was no significant difference in the levels at 4, 7, 9, or 10 weeks of gestation between the groups (Table 1).

**Table1.** Serum E2 levels over time in the non-HR and HR groups

|  | non-HR group (n) | HR group (n) | P value |
| --- | --- | --- | --- |
| 4 weeks | $453.8\pm137.7$ (5) | $291.9\pm200.7$ (75) | *NS* |
| 5 weeks | $622.3\pm379.7$ (15) | $351.3\pm173.0$ (75) | $<0.05$ |
| 6 weeks | $870.1\pm326.7$ (12) | $644.1\pm262.3$ (74) | $<0.01$ |
| 7 weeks | $2056.3\pm1635.0$ (3) | $976.6\pm342.2$ (73) | $<0.01$ |
| 8 weeks | $3232.0\pm1781.3$ (10) | $1285.7\pm453.4$ (71) | $<0.01$ |
| 9 weeks | $1589.3\pm660.9$ (32) | $1556.3\pm587.9$ (62) | *NS* |
| 10 weeks | $1790.9\pm974.6$ (42) | $1781\pm649.2$ (36) | *NS* |

Data are expressed as mean ± SD, NS; not significant, unit: pg/mL

The serum P4 values in the non-HR and HR groups were 23.9±5.1 and 14.0±7.8 ng/mL at 4 weeks, 25.8±12.7 and 14.7±8.6 ng/mL at 5 weeks, 22.7±326.7 and 15.4±7.5 ng/mL at 6 weeks, 28.9±18.0 and 18.8±7.6 ng/mL at 7 weeks, 30.9±14.1 and 22.4±7.4 ng/mL at 8 weeks, 25.8±7.2 and 26.9±8.9 ng/mL at 9 weeks, and 28.5±8.8 and 27.6±9.3 ng/mL at 10 weeks of gestation, respectively. The serum P4 level was significantly higher in the non-HR group than in the HR group at 4, 5, and 6 weeks of gestation (4 weeks, p <0.01; 5 weeks, p <0.01; 6 weeks, p <0.01). There was no significant difference in the levels at 7, 8, 9, or 10 weeks of gestation between the groups (Table 2).

**Table2.** Serum P4 levels over time in the non-HR and HR groups

|  | non-HR group (n) | HR group (n) | P value |
| --- | --- | --- | --- |
| 4 weeks | 23.9±5.1 (5) | 14.0±7.8 (75) | <0.01 |
| 5 weeks | 25.8±12.7 (15) | 14.7±8.6 (75) | <0.01 |
| 6 weeks | 22.7±326.7 (12) | 15.4±7.5 (74) | <0.01 |
| 7 weeks | 28.9±18.0 (3) | 18.8±7.6 (73) | NS |
| 8 weeks | 30.9±14.1 (10) | 22.4±7.4 (71) | NS |
| 9 weeks | 25.8±7.2 (32) | 26.9±8.9 (62) | NS |
| 10 weeks | 28.5±8.8 (42) | 27.6±9.3 (36) | NS |

Data are expressed as mean ± SD, NS; not significant, unit: ng/mL

## Discussion

Although some studies have reported that the use of ART in breast cancer survivors does not significantly affect the prognosis of cancer [11] [12] [13] [14] [15], the number of such studies is small, and the safety of ART has not been established to date. Previous reports have indicated that if women without breast cancer receive more than six sessions of controlled ovarian stimulation (COS) under ART, the standardized incidence rate of breast cancer increases 1.23-times [16]. Exposure to estrogen causes dose-dependent cellular proliferation and an increase in cancer cell lines estrogen receptor [17]. Hence, careful consideration is necessary before offering this option to patients with cancer. As serum estrogen levels increase with COS, the risk of recurrence is of particular concern in patients with hormone receptor-positive breast cancer[18].

Estrogen and its metabolites have been implicated in the onset and progression of breast cancer [19]. Recently, COS using gonadotropin-releasing hormone (GnRH) antagonists and aromatase inhibitors was reported to be effective for minimizing any increase in the serum E2 levels [20][21]. However, when aromatase inhibitors are used for COS, the serum P4 level is maintained at a comparatively high level [22]. The use of a GnRH agonist rather than of hCG as the trigger may be effective for minimizing the increase in the estrogen and progesterone levels [23] [24] [25].

Conventionally, progesterone levels do not affect the risk of breast cancer [26], although some studies have reported that progesterone levels may contribute toward breast cancer risk [27] [28]. Therefore, a method in which the serum E2 and P4 levels do not increase above the required levels must be selected when using ART for breast cancer survivors.

In ART, HRT is important for creating the structural and functional environment for embryo implantation in the endometrium and is an established treatment with proven efficacy [29]. With respect to the hormones administered, progesterone and a combination of progesterone and estrogen have been reported to be effective [29] [30]. In a fresh embryo transfer, the GnRH agonist used for COS as a short or long protocol and the GnRH antagonist used for pituitary suppression prevent adequate corpus luteum formation, causing luteal function failure [31] [32]. This indicates that HRT with estrogen and progesterone is required for both implantation and continuation of the pregnancy. In a frozen-thawed embryo transfer and during HRT, ovulation does not occur; thus, corpus luteum is not formed and progesterone is not secreted endogenously. If the corpus luteum is removed before 7 weeks of gestation, the serum P4 level suddenly decreases and miscarriage may occur [33]. Thus, luteal support is necessary until at least 7 weeks of gestation. During the natural cycle, luteal support is not necessarily required; however, it is needed if there are luteal phase defects. Many women with decreased ovarian function after neoadjuvant chemotherapy for breast cancer who do not have an ovulatory cycle and whose fertility has declined are likely to require HRT, including the administration of estrogen and progesterone. This plays a major role in embryo implantation and the continuation of pregnancy.

Similar to COS, estrogen and progesterone replacement therapies are administered to patients without an ovulatory cycle who are undergoing thawed embryo transfer. The effect of these therapies on breast cancer is concerning. Currently, the safety of these replacements is not known. In this study, we measured the levels of serum E2 and P4 levels in the first trimester of pregnancy in patients who did not receive HRT.

Our results suggest that in the first trimester, the serum E2 and P4 levels in the HR group were not higher than those in the non-HR group. Thus, the amount of HRT required to establish a pregnancy may not increase the cancer risk associated with hormonal exposure in patients with hormone receptor-positive breast cancer compared to that in those with spontaneous pregnancy.

We found that the serum E2 levels were significantly lower in the HR group than in the non-HR group at 5, 6, and 8 weeks of gestation. Differences at 4 and 7 weeks of gestation may not have been statistically significant owing to the small sample size. In the non-HR group, the levels tended to increase over time (Table 1). At approximately 6–7 weeks of gestation, the main site of hormone production switches from the corpus luteum to the placenta (luteo-placental shift) [34], and the levels rapidly increased during this period (Fig 4a). There was no significant difference in the serum E2 levels at 9 and 10 weeks of gestation between the two groups; this may be because estrogen replacement therapy was discontinued in the HR group at 8 weeks of gestation, and in both groups, the main site of hormone production was now the placenta.

The serum P4 levels were significantly lower in the HR group than in the non-HR group at 4, 5, and 6 weeks of gestation (Table 2). In the non-HR group, the serum P4 levels gradually increased between 4 and 10 weeks of gestation to 20–30 ng/mL (Fig 4b), and in the HR group, the level remained at approximately 15 ng/ml, which was significantly lower than that in the non-HR group. As these pregnancies were achieved by embryo transfer and HRT with estrogen and progesterone, without the formation of the corpus luteum, the serum P4 levels would have been solely derived from the transvaginal natural progesterone administered. Although the use of transvaginal natural progesterone results in significantly lower serum P4 levels than those achieved after intramuscular injection of natural luteal hormone, the P4 concentration in the endometrial tissue is significantly higher; hence, the serum P4 level does not reflect the local concentration in the tissue [35] [36]. In practice, the serum P4 levels are reported to be low after the administration of transvaginal natural progesterone, but the P4 concentration in the endometrial tissue is maintained at a higher level [37]. In this study, because few samples were collected from patients in the non-HR group at 7 weeks of gestation, we could not evaluate this aspect. However, as the placenta started to produce hormones by approximately 6–7 weeks of gestation, the hormone levels increased after 8 weeks of gestation in the HR group, and the significant difference between the groups disappeared.

Our results suggest that increased serum E2 and P4 levels in the first trimester of pregnancy owing to HRT are lower than the serum E2 and P4 levels noted during the first trimester of spontaneous pregnancy or pregnancy following regular infertility treatment. This suggests that the risk of breast cancer associated with thawed embryo transfer with HRT of estrogen and progesterone may not be greater than that associated with spontaneous pregnancy.

This study has several limitations. First, only a few samples were collected from the patients in the non-HR group. Therefore, we contained patients undergoing medication for ovulation induction in non-HR group. This was unavoidable because the study design involved the use of left-over blood from blood drawn during scheduled hospital visits by women who conceived naturally or after IUI. Second, because this study did not include breast cancer survivors, we were unable to assess the prognosis or risk of recurrence in breast cancer survivors who underwent thawed embryo transfer with HRT of estrogen and progesterone. It is an ethical dilemma not to offer HR to breast cancer survivors when they have a rare opportunity to conceive using ART. Hence, we opted for a study design in which we compared the non-HR and HR groups.

In conclusion, our results showed that thawed embryo transfer with HRT did not increase the serum hormone levels beyond those observed in spontaneous pregnancy. This suggests that in patients with hormone receptor-positive breast cancer, HRT administered during the first trimester of a pregnancy, established as a result of ART, after treatment of the primary disease may not increase the sex hormones levels beyond those observed in spontaneous pregnancy.

## Acknowledgements

The authors would like to thank to all colleague of the Department of Obstetrics and Gynecology, Reproduction Center, Department of Laboratory Medicine, Toho University Omori Medical Center for the cooperation of the subjects’ recruitment and measuring blood samples.

